# The effects of breeding timing, clutch size, and nesting strategy on reproductive success in the crested ibis (*Nipponia nippon*)

**DOI:** 10.64898/2026.02.04.703896

**Authors:** Yuansi He, Xuebo Xi, Siyi Zeng, Ke Wang, Xiang-Yi Li Richter, Daiping Wang

**Affiliations:** Institute of Zoology, Chinese Academy of Sciences; Administration Bureau of Dongzhai National Nature Reserve, Luoshan, Henan Province; University of Bern

**Keywords:** Crested ibis (*Nipponia nippon*), breeding timing, clutch size, nesting strategies, reproductive success

## Abstract

1. Studying the reproduction process, which is a key determinant of individual and population fitness in endangered species, is challenging but urgently needed. The crested ibis (*Nipponia nippon*), a flagship endangered species recovering from an extreme population bottleneck, provides a valuable opportunity to examine how life-history strategies shape reproductive success and inform future conservation practices.
2. We monitored 176 breeding pairs of crested ibis over three consecutive breeding seasons and investigated the effects of three key life-history traits, namely breeding timing, clutch size, and nesting strategy (solitary versus colonial), on reproductive success (hatching and fledging success).
3. Our analysis found that both hatching and fledging success declined significantly as breeding initiated later, and a positive association between clutch size and reproductive success in this species. These patterns were robust and repeatable across three years. Unlike other closely related species in this family, sibling competition is generally non-lethal, leading to large clutch sizes fledged in this endangered species. We consider this pattern to be a main reason underlying the rapid population recovery observed in the crested ibis. On the other hand, nesting strategy (colonial vs. solitary breeding) had no detectable effect on reproductive success. This pattern indicates the crested ibis can adopt different breeding strategies across habitats, highlighting its capacity to flexibly adjust breeding behavior in response to local environmental conditions.
4. Our results provide an integrative assessment of how key life-history traits shape reproductive outcomes in a wild population of the crested ibis, serving as a foundation for evaluating its current status of population recovery and refining future conservation strategies for endangered avian species sharing similar life-history characteristics.

## Introduction

Reproduction is a fundamental component of a species’ life history because it directly determines individual fitness while imposing substantial energetic and physiological costs. As resources are inherently limited, individuals must allocate energy among competing physiological and behavioral demands in ways that are worth the cost and ultimately maximize fitness (Williams, 1966; Bell, 1980; Stearns, 1989). For long-lived species capable of reproducing multiple times over their lifespan, increased reproductive effort in a given breeding attempt may compromise future survival, reproductive output, or offspring quality, a phenomenon widely referred to as the cost of reproduction (Perrins, 1965; Williams, 1966; Hanssen et al., 2005). Thus, individuals are selected to regulate their reproductive effort in each breeding opportunity. In birds, especially those living in seasonal environments, the most important reproductive decisions they have to make include the timing of breeding (laying date) and reproductive effort (clutch size) (Daan & Tinbergen, 1997; Houston & McNamara, 1999). Birds must synchronize breeding with periods of favorable environmental conditions while simultaneously adjusting clutch size to the maximum number of offspring they can successfully raise, so as to reach an optimal level that yields maximal fitness (Martin, 1987; Rowe et al., 1994; Reséndiz-Infante & Gauthier, 2020). Beyond these temporal and clutch-size decisions, birds also face choices regarding the spatial and social context of reproduction (nest solitarily or in colonies). Multiple interacting intrinsic and extrinsic factors can influence the relative benefits of these strategies, resulting in different optimal strategies under varying ecological and social conditions (Hamilton & May, 1977; Götmark & Andersson, 1984; Neil J. Buckley, 1997; Tella et al., 2001; Hoogland, 2013; Minias et al., 2015).

Breeding timing has long been recognized as a major determinant of avian reproductive success. Many bird species exhibit a consistent seasonal decline in reproductive performance. In single-brooded species, reproductive success often decreases linearly across the season, whereas in multi-brooded species, there is usually an optimum in the middle of the season, followed by a steady decline (Verhulst & Nilsson, 2008). It may either be a consequence of timing per se, affecting all individuals in the same way (the ‘date’ hypothesis), or reflect the quality differences between breeders, referring to the phenotypic condition of the breeding individuals and/or their territories

(the ‘quality’ hypothesis) (Christians et al., 2001; Verhulst & Nilsson, 2008; Karagicheva et al., 2016). Under the ‘date’ hypothesis, the decline of reproductive success may be due to some factors that deteriorates with season, which may include 1) absolute date (e.g. the need for reserved time and energy for post-breeding activities like molting, wintering and migration or seasonal change of weather) (Nilsson & Svensson, 1996; Olmos, 2003), 2) relative date (e.g. chicks fledging earlier survive better than those fledging later during the post fledging period) (Nilsson & Smith, 1988; Verhulst & Tinbergen, 1991), 3) food availability (e.g. early breeders better match the shifting phenology of underlying levels of the food chain) (Reed et al., 2013) and 4) predation pressure (e.g. breeding early may reduce exposure to periods of peak predator activity) (Götmark, 2002). While the ‘quality’ hypothesis posits that early breeders are intrinsically superior, either in phenotypic condition, territory quality, or both, and therefore achieve higher reproductive success (Christians et al., 2001; Liu et al., 2025). Nonetheless, both effects are expected to be prevalent, because only when date per se matters does early breeding confer a selective advantage to high-quality individuals (Verhulst & Nilsson, 2008).

Clutch size represents another fundamental life-history trait in birds. Lack (1947, 1954) first proposed that natural selection has adjusted the clutch size to the largest number of young for which the parents can rear on average. Klomp (1970) further refined this idea, stating that natural selection favors an optimal and most productive clutch size, which maximizes the number of offspring that survive to independence and contribute to future reproduction. It is now widely accepted that clutch size is shaped not only by environmental resources like food (Lack, 1947; Martin, 1987; Rowe et al., 1994), but also by trade-off between investment in current reproduction and future fitness costs, as well as the allocation of parental effort among offspring (Williams, 1966; Charnov & Krebs, 1974; Bell, 1980; Siikamäki, 1998; Stearns, 1998). Within this framework, clutch size not only determines the number of offspring produced, but also influences the competitive environment experienced by nestlings. As clutch size increases, sibling competition becomes inevitable. In birds, younger chicks are often less competitive and thus more prone to being suppressed by older chicks in food-seeking, and sometimes die from starvation. In species exhibiting severe or obligate siblicide like black eagle (*Aquila verreauxi*) and masked booby (*Sula dactylatra*), the youngest chick is frequently killed by older siblings even when food resources are sufficient (Gargett, 1977; Mock, 1984). Despite the apparent wastefulness of producing offspring that may not survive, extensive research has demonstrated that later-hatched nestlings can confer adaptive benefits under certain conditions. These include providing “extra reproductive value” when additional offspring can be successfully reared, and/or serving as “insurance reproductive value” by replacing an older sibling that dies early (Mock et al., 1990). In the former occasion, parents may produce more offspring than they can typically rear as a hedge against environmental uncertainty. When food resources become limiting, weaker offspring are eliminated early to reduce parental costs and allow greater investment in the remaining young. Conversely, when food is abundant, the full brood may be raised, maximizing reproductive output (Lack, 1954; Husby, 1986; Mock & Forbes, 1994). In the latter situation, which usually occurs in species practicing obligate siblicide, younger chicks function primarily as an insurance against the early loss or infirmity of older chicks. If the older chick is unable to eliminate the younger, dominance may be reversed, resulting in the survival of the initially subordinate offspring (Gargett, 1977; Cash & Evans, 1986; Mock et al., 1990). The outcomes of sibling competition are further shaped by parental strategies. In addition to allowing competitive interactions to proceed without intervention, parents may actively regulate the intensity of within-brood competition by adjusting feeding preferences, physically preventing aggressive interactions, or diverting the attention of dominant chicks (Leonard et al., 1988; Wiebe & Bortolotti, 2000; Szojka et al., 2020; Brode et al., 2021). Collectively, these processes indicate that the clutch size maximizing reproductive returns varies among species and even among breeding attempts within the same species, as it is influenced by multiple interacting factors, including environmental conditions, individual quality, life-history trade-offs, and offspring competitive dynamics.

In addition to decisions about the timing and investment intensity of breeding, individuals face fundamental choices about where and with whom to breed. Whether to nest solitarily or in colonies represents a key reproductive and spatial decision, yet no single strategy is universally optimal across all ecological and social contexts. Natural selection may favor a diversity of strategies that balance competing selective pressures, resulting in a mixed evolutionarily stable strategy (ESS) within populations (Smith, 1973, 1982). Intrinsic factors, particularly those related to kinship and social structure, can impose fundamental constraints shaping breeding strategies. Theoretical models have shown that competition among close relatives may favor dispersal even in stable environments, thereby influencing whether individuals breed in isolation or in close proximity to conspecifics (Hamilton & May, 1977). While cooperation among kin may prompt spatial aggregation and the formation of breeding groups. Empirical evidence from prairie dogs (*Cynomys ludovicianus, C. gunnisoni* and *C. parvidens*) shows that it is the absence of close kin that highly raises the possibility of young females dispersing from their natal sites, highlighting the role of kinship in dispersal and breeding-site decisions (Hoogland, 2013). In parallel, extrinsic ecological factors, including predation pressure, resource availability, and parasite load, can modulate the fitness consequences of different breeding strategies, altering the cost–benefit balance of breeding solitarily versus colonially. High predation risk may favor colonial breeding through mechanisms such as predator dilution, enhanced vigilance, and communal defense, whereas abundant resources can alleviate intra-group competition and facilitate aggregation (Andersson & Wiklund, 1978; Götmark & Andersson, 1984; Murphy & Schauer, 1996; Roberts, 1996; Neil J. Buckley, 1997; Minias et al., 2015). Conversely, increased competition for food, poor physical condition caused by social stress and elevated parasite load will selects for dispersed nesting, particularly under conditions of limited resources or high breeding density (Brown et al., 2001; Tella et al., 2001). Together, multiple factors and selective forces shape animals’ decisions regarding breeding location and social context. Diversity and flexibility in breeding strategies can be interpreted as an adaptive form of bet-hedging in response to variable ecological and social conditions, allowing solitary and colonial breeding to persist as context-dependent, evolutionarily stable outcomes within populations (Slatkin, 1974).

The crested ibis (*Nipponia nippon*) is one of the most famous endangered bird species that has recovered from an extremely small population (Liu, 1981; Li, 2023). Historically widespread across Northeast Asia, (BirdLife-International, 2001; Li et al., 2014), the species suffered a dramatic population decline during the twentieth century due to habitat destruction and intensive human activities (e.g. illegal hunting, overuse of fertilizer and pesticide) (Archibald et al., 1980; Ding, 2004; Li et al., 2009; Feng et al., 2019). Until 1981, a small wild population of seven individuals, including two breeding pairs and three chicks belonging to one of the couples, was rediscovered in Yangxian County of Shaanxi Province, China (Liu, 1981). Since then, extensive conservation and reintroduction programs have been implemented to save this endangered species, and so far, the global population of wild and captive crested ibis has been thought to exceed 10,000 (Li, 2023), with several populations reintroduced to its historical ranges in China (Yu et al., 2015; Zheng et al., 2018; Xie et al., 2020; Qiu et al., 2023; Cai et al., 2024), Korea (Choi et al., 2020), and Japan (Wajiki et al., 2014; Okahisa et al., 2022). As a long-lived species, the crested ibis can breed repeatedly throughout its lifetime (Shi & Cao, 2001; Yu et al., 2007), with each breeding attempt requiring an extended period of incubation and parental care, such that typically only one brood can be successfully raised per breeding season (Shi et al., 1999; Zhai et al., 1999). These life-history characteristics imply that adult individuals must balance investment between current and future reproductive opportunities, and carefully adjust breeding timing and clutch size within a limited breeding window. Also, as an altricial species, crested ibis chicks exhibit pronounced developmental asymmetries within broods. Such developmental differences often promote sibling aggression, and increasing clutch size can be associated with intensified intra-brood competition for parental resources. However, in contrast to many other species in Threskiornithidae, pecking among the chicks of the crested ibis is considered to be largely ritualized and non-lethal (Li et al., 2004). Consequently, larger clutches may not be associated with markedly intensified intra-brood competition and do not necessarily result in reduced reproductive success in this species. Moreover, previous research has shown that the breeding strategy (colonial versus solitary nesting) of crested ibis varies across space and time (Shi & Cao, 2001; Liu et al., 2003), suggesting a capacity for flexible adjustment of reproductive strategies in response to local environmental conditions. To date, substantial knowledge has been accumulated on breeding, growth and development in captivity, release practices, basic reproductive indicators, population dynamics, and habitat use in reintroduced crested ibis populations (Endo & Nagata, 2013; Yu et al., 2015; Dong et al., 2018; Liu et al., 2020; Li et al., 2021; Qiu et al., 2023; Cai et al., 2024; Xi et al., 2025; He, Xi, et al., 2025; He, Zhang, et al., 2025). However, studies explicitly quantifying these key fundamental life-history factors influencing reproductive success in wild populations remain relatively scarce. In addition, systematic investigations of reintroduced populations outside Shaanxi Province are still limited. To this end, examining how major life-history traits shape reproductive success in endangered species, particularly flagship species like crested ibis, is essential for effective conservation and for accurately predicting population dynamics.

Here, we investigated the effects of fundamental life-history traits on reproductive success in a wild population of the crested ibis. Specifically, based on the environmental conditions of the reserve and the species’ life-history traits, we examined how breeding timing, clutch size, and nesting strategy (solitary versus colonial) were associated with hatching success and fledging success over three consecutive breeding seasons (2023∼2025). By testing these relationships, our study aims to provide new insights into the reproductive ecology of the crested ibis, improve understanding of factors influencing population recovery, and offer empirical evidence to inform the development of targeted conservation strategies and the evaluation of ongoing reintroduction efforts.

## Methods

### Study species

The crested ibis is a socially monogamous species characterized by high mate and nest-site fidelity, with a maximum recorded lifespan of 17 years in the wild (Shi & Cao, 2001; Yu et al., 2007). The breeding season typically begins in March, during which both members of a breeding pair participate in all stages of reproduction, including nest building, incubation, and chick rearing (Zhai et al., 2011). Crested ibises usually nest in tall trees. The mean height of nesting trees is approximately 16.1 m, and nests are typically located around a height of 12.6 m above the ground (Xi et al., 2025). The species usually produces one clutch per breeding season, with an interval of one to two days between egg-laying and a clutch size ranging from two to four eggs (Shi et al., 1999; Ding, 2004). The incubation period lasts approximately 28 days, followed by a nestling period of 40–45 days (Shi et al., 1999; Zhai et al., 1999). After fledging, juveniles remain dependent on their parents for a period of time while learning essential survival skills, such as foraging (Ding, 2004). Early studies reported that the crested ibis nested solitarily and exhibited territoriality in the breeding season (Shi & Cao, 2001). However, with ongoing population recovery and range expansion, an increasing tendency toward colonial nesting has been documented in recent decades (Liu et al., 2020).

### Study area and population background

The study was conducted in Dongzhai National Nature Reserve (31°58’ N, 114°17’ E) and surrounding areas in Henan Province, China, covering a total area of approximately 73,000 ha. The region lies within the transitional zone between the northern subtropical and warm temperate climates and is characterized by a warm, humid climate with four distinct seasons. Mean annual temperature is 15.1 °C, and mean annual precipitation is approximately 1,200 mm. The topography gradually descends from southwest to northeast and includes mountainous, hilly, ridge, and plain landscapes, with elevations ranging from 100 to 840 m above sea level. The mosaic of forest patches and extensive rice paddies provides a suitable foraging and nesting habitat for crested ibises.

In 2007, 17 crested ibises were introduced to Dongzhai National Nature Reserve to establish a captive breeding population. The captive population subsequently increased, and a reintroduction program was initiated in 2013. Between 2013 and 2023, 135 individuals were released into the wild during seven release events (34 in 2013, 28 in 2014, 18 in 2015, 22 in 2017, 20 in 2021, 7 in 2022, and 6 in 2023). These efforts successfully established a self-sustaining reintroduced wild population (Xi et al., 2025).

### Nest searching and monitoring

During the breeding seasons from 2023 to 2025 (February to June each year), we conducted systematic monitoring of crested ibis breeding activities throughout the study area. Nest searches initially focused on historical breeding sites by checking for signs of ibis activity, such as fecal traces and newly added nesting materials. Potential nesting sites were further identified using multiple information sources: (1) nocturnal roosting locations derived from GPS transmitters deployed on a subset of individuals, (2) eyewitness observation records of adult birds with breeding plumage during field surveys, and (3) information on the locations of breeding nests provided by local residents.

Once a nest was located, systematic follow-up monitoring was initiated to comprehensively document the reproductive progress and final fate. During the nest-building stage, to minimize human disturbance as much as possible, monitoring was conducted at a frequency of one to two visits per week. Observations were made from a distance using binoculars, with detailed records of nest construction progress and the behavioral patterns of adult birds.

The crested ibis is an asynchronously hatching species and incubation commences after the first egg is laid (Ding, 2004). Accordingly, the first observation of a stable incubation posture was used to indicate that egg laying had begun no later than that date, and the monitoring frequency was increased to once every two days to document incubation status and any irregularities (e.g., egg loss). To determine the exact first hatch date, the monitoring frequency increased to daily as the expected hatching date approached. Hatching was confirmed by the concurrent observation of eggshell fragments beneath the nest and the initiation of parental chick-rearing behavior. For nests found after hatching, we estimated the first hatch date from chick body size, typically limiting the error to 1–3 days. Finally, the first egg-laying date was derived by back-calculating 28 days (the incubation period) from the confirmed first hatch date.

The clutch size and hatching success were determined by combining earlier monitoring records with counts of chicks and unhatched eggs obtained from real-time drone imagery (DJI Mini 4 Pro; DJI, Shenzhen, China). This drone survey was conducted within one week post-hatching, taking care to avoid disturbing the attending parents. For nests with larger clutch sizes, where hatching intervals among eggs may be relatively long, multiple observations were required to determine the final number of successfully hatched chicks. During the nestling period, monitoring was conducted once every two days, focusing on chick growth, development, and survival dynamics (Xi et al., 2025).

At 25–30 days of age, each chick was fitted with a unique combination of a metal leg band (issued by the National Bird Banding Center of China) and a colored plastic band for individual identification and subsequent field tracking. Concurrently, we collected standardized morphometric data and biological samples from each individual. Monitoring continued post-banding until fledging to determine the final number of fledglings.

In this study, we have monitored 176 nesting attempts over three years (45 in 2023, 64 in 2024, and 67 in 2025).

### Definition of reproductive performance and nesting type

In this study, the first egg-laying date was used as a proxy for breeding timing. To standardize timing across years, we defined March 1 as Day 1 of the breeding season. All subsequent dates were then converted to a ‘Day of Season’ value, calculated as the calendar date count starting from March 1. For example, March 1 = 1, March 2 = 2, and April 15 = 46.

Based on field records of clutch size, the number of hatched eggs, and the number of fledglings, we quantified reproductive performance for each nest by calculating hatching success (number of hatched eggs/clutch size) and fledging success (number of fledglings/clutch size).

Nests were categorized as either colonial or solitary. The study area comprises primarily small forest patches dominated by Masson pine (*Pinus massoniana*) or loblolly pine (*Pinus taeda*), embedded within a farmland and village matrix. These patches had a mean area of 1.48 ± 0.95 ha (mean ± SD; n = 43). Consequently, a nest was classified as colonial if ≥2 breeding pairs nested within the same forest patch during the same breeding season; otherwise, it was classified as solitary. Over the three-year study, we identified 72 colonial nests (20 in 2023, 28 in 2024, 24 in 2025) and 104 solitary nests. Colony size ranged from 2 to 10 breeding pairs, averaging 3.4 ± 2.2 pairs (n = 21 colonies; Table S1). The mean nearest-neighbor distance was 24.27 ± 20.52 m for colonial nests (n = 72), which was significantly less than that for solitary nests (2581.66 ± 2995.42 m; n = 104).

### Statistical analysis

We examined annual variation in reproductive performance and the effects of breeding timing, clutch size, and nesting type on reproductive success using a combination of linear models and linear mixed-effects models. Sample sizes vary across models because nests lacking confirmed hatching dates or reliable clutch size data were excluded from the corresponding analyses. All analyses were performed in R (R Core Team, 2022). Linear mixed-effects models were fitted using the lme4 package (Bates et al., 2015). Inference on fixed effects in mixed-effects models was conducted using the lmerTest package, which provides approximate degrees of freedom and two-sided P values based on Satterthwaite’s method (Kuznetsova et al., 2017). Statistical significance was evaluated at α = 0.05.

We first assessed inter-annual variation of reproductive success in this species (i.e., hatching success rate and fledging success rate). To this end, we used two linear models (Model 1 and 2) with the hatching success rate and fledging success rate as the response variable, respectively. The ‘year’ (a categorical variable with three levels) was added as the fixed effect.

To evaluate the effect of breeding timing on reproductive performance across years, we fitted another two linear mixed-effects models with hatching success rate and fledging success rate as the response variable, respectively (Model 3 and 4). The breeding timing (i.e., the first egg-laying date) was added as the fixed effect. In addition, the ‘year’ was added as the random effect to account for unmeasured interannual heterogeneity. Importantly, to further assess whether the effect of breeding timing on reproductive performance was consistent across three years, we conducted additional separate linear models for each breeding season (2023, 2024, and 2025: Model 5, Model 6 and Model 7 for the hatching success; Model 8, Model 9 and Model 10 for the fledging success). In these linear models, hatching success rate and fledging success rate were analyzed separately as response variables, with breeding timing as the explanatory variable.

Next, to test whether the effect of reproductive investment (i.e., clutch size) on reproductive success (i.e., hatching success and fledging success rate), we fitted another two linear mixed-effects models with the hatching success rate and fledging success rate as the response variable, respectively (Model 11 and 12). The ‘clutch size’ was added as the fixed effect and the ‘year’ was added as the random effect to account for unmeasured interannual heterogeneity.

Finally, we assessed the effect of nesting type (colonial vs. solitary) on reproductive success by fitting separate linear mixed-effects models for hatching success and fledging success rate, respectively (Model 13 and 14). Each model included nesting type (a categorical variable with two levels) as the fixed effect and year as the random intercept.

## Results

### Overview of reproductive success

A total of 176 nesting attempts were monitored from 2023 to 2025 (45 in 2023, 64 in 2024, and 67 in 2025). Of those, 160 attempts resulted in successful nest construction and produced a total of 502 eggs, of which 340 hatched into chicks. Eventually, 236 of these chicks fledged (see Table S2 for detailed information). The average hatching success rate was 0.652 ± 0.030 (Mean ± SE), and the average fledging success rate was 0.460 ± 0.032 (Mean ± SE) (Table 1). The results of the linear model showed that there was no significant difference in the hatching success rate between 2024 and 2023, nor between 2025 and 2023. The fledging success rate in 2024 was significantly higher than that in 2023 (*P* < 0.01), while there was no significant difference in the fledging success rate between 2025 and 2023 (Table 1).

**Table 1.** Variation of reproductive success in the crested ibis in Dongzhai in 2023–2025 (Model 1 and 2). Annual hatching/fledging success rates are reported as the mean ± SE of per-nest proportions.

|  | Hatching success rate | Difference<br>(vs. 2023) | Fledging success rate | Difference<br>(vs. 2023) |
| --- | --- | --- | --- | --- |
| 2023 | $0.592 \pm 0.056$ | - | $0.320 \pm 0.055$ | - |
| 2024 | $0.701 \pm 0.049$ | $t = 1.327$<br>$P = 0.187$ | $0.571 \pm 0.055$ | $t = 2.899$<br>**$P = 0.004$** |
| 2025 | $0.649 \pm 0.051$ | $t = 0.701$<br>$P = 0.484$ | $0.448 \pm 0.049$ | $t = 1.501$<br>$P = 0.135$ |
| Overall | $0.652 \pm 0.030$ | - | $0.460 \pm 0.032$ | - |

### Effects of breeding timing on reproductive performance

We found that breeding timing had a strong, significant effect on both hatching and fledging success in our study (Figures 1 and 2). Specifically, hatching success rate declined significantly with later breeding initiation, indicating that nests established later in the season experienced reduced egg hatching success (*β* ± SE = -0.016 ± 0.003, *t* = -5.217, *P* < 0.001, n = 145, Figure 1a; Table S3, Model 3). Importantly, we found this pattern was robust when we separated the dataset per year, and repeatable across three years (Figure 1b, c, d; Table S4, Model 5, Model 6 and Model 7). Similarly, fledging success rate also decreased significantly with later breeding timing, suggesting that the negative effects of delayed breeding persisted beyond the hatching stage and influenced offspring survival through to fledging (*β* ± SE = -0.015 ± 0.003, *t* = -4.380, *P* < 0.001, n = 144, Figure 2a, Table S5, Model 4). Again, this pattern is robust within each year and repeatable across three years (Figure 2b, c, d; Table S6, Model 8, Model 9 and Model 10). Detailed results of the mixed-effects and year-specific models are provided in Tables S3–S6.

**Figure 1.**
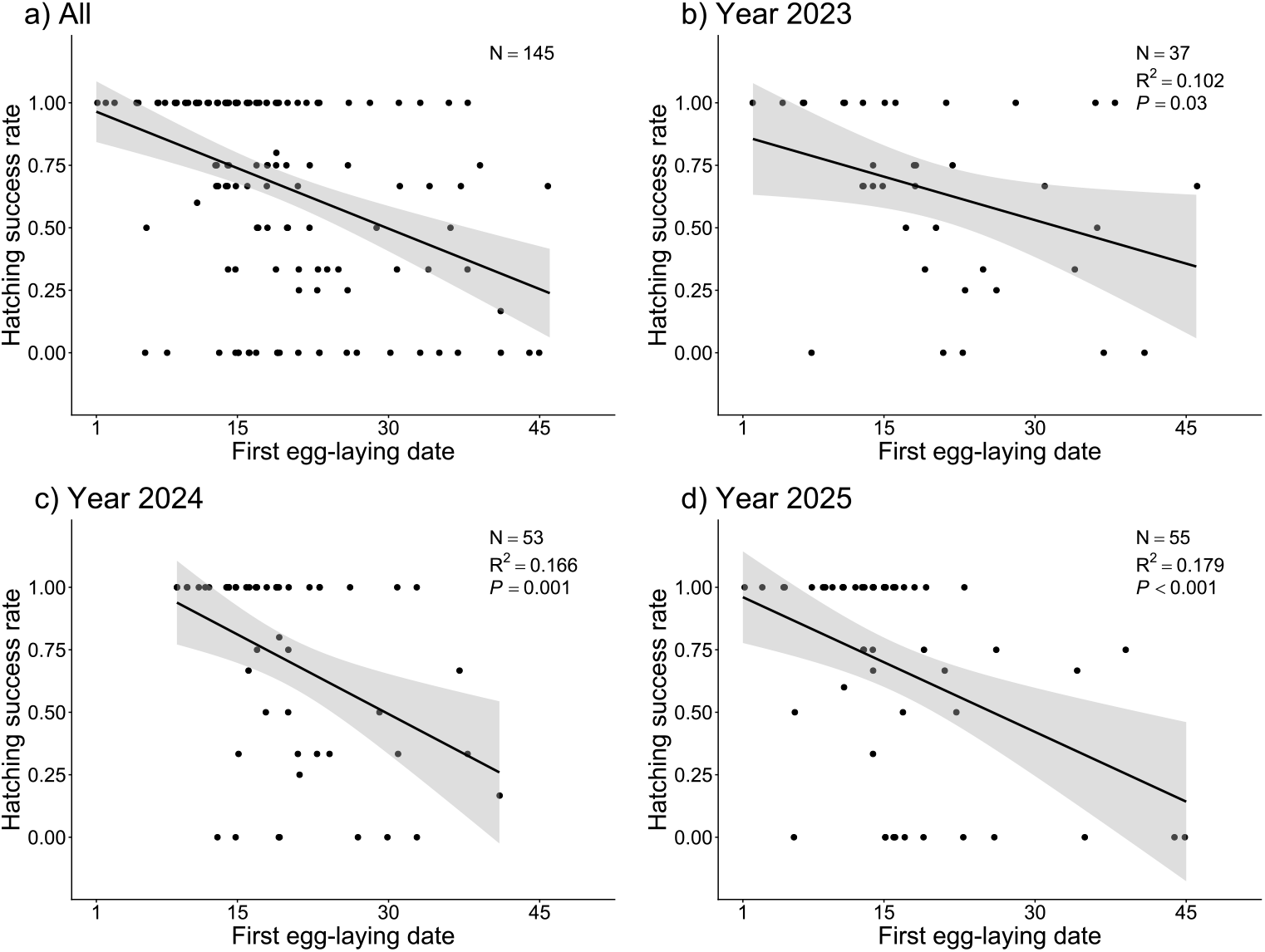
Relationship between hatching success rate and the first egg-laying date. Panel (a) shows the combined dataset across all three years (2023–2025; Model 3, Table S3), while panels (b–d) show data for 2023 (Model 5, Table S4), 2024 (Model 6, Table S4), and 2025 (Model 7, Table S4), respectively. Fitted lines and confidence intervals are shown for visualization only and were obtained using simple linear regression.

**Figure 2.**
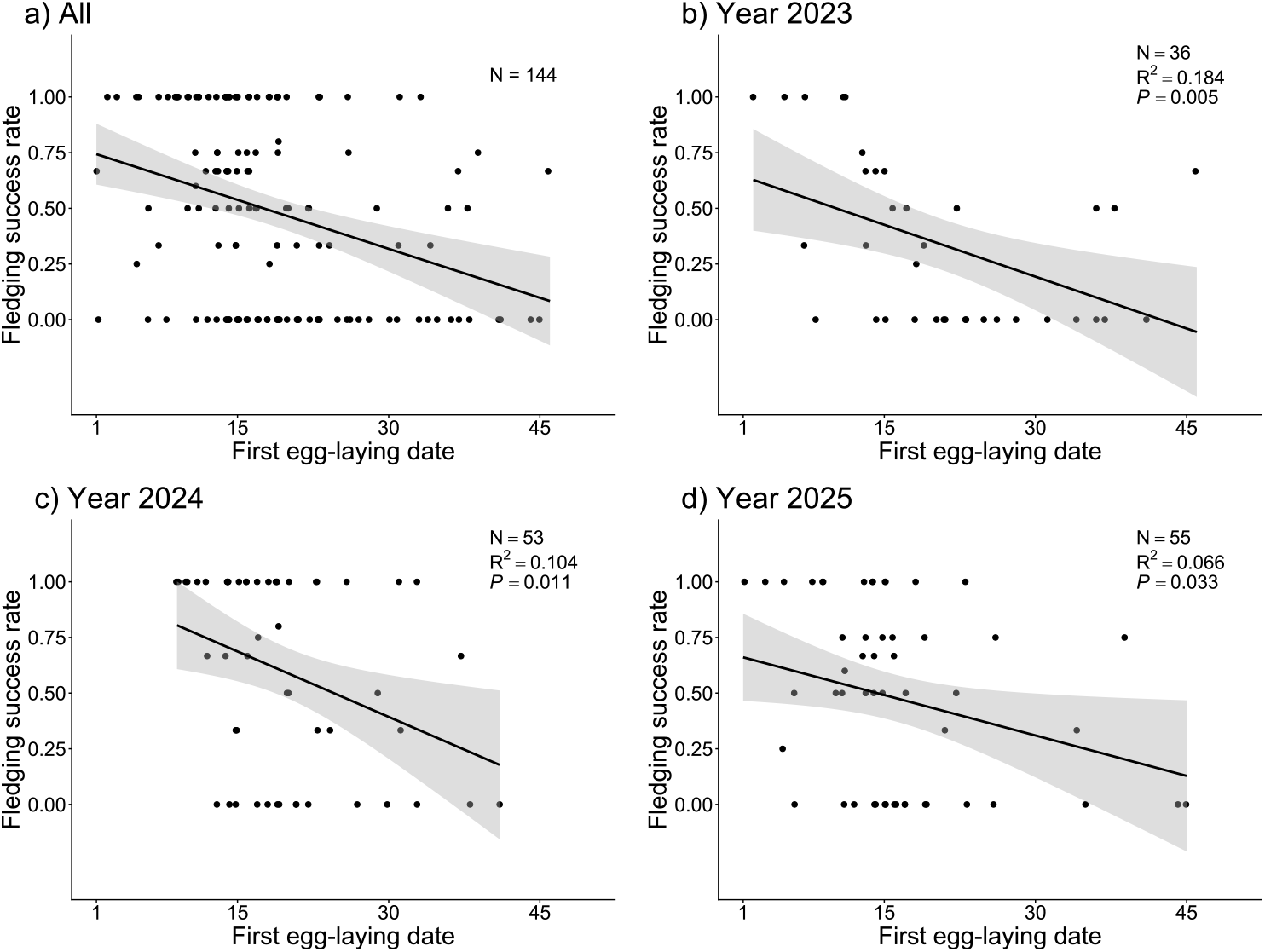
Relationship between fledging success rate and the first egg-laying date. Panel (a) shows the combined dataset across all three years (2023–2025; Model 4, Table S5), while panels (b–d) show data for 2023 (Model 8, Table S6), 2024 (Model 9, Table S6), and 2025 (Model 10, Table S6), respectively. Fitted lines and confidence intervals are shown for visualization only and were obtained using simple linear regression.

### Effects of clutch size on reproductive success

Clutch size was positively associated with reproductive success in our study. Specifically, hatching success rate increased significantly with larger clutch size, indicating that nests with more eggs achieved higher egg hatching success (*β* ± SE = 0.141 ± 0.032, *t* = 4.447, *P* < 0.001, n = 154; Figure 3a; Table S7, Model 11). Similarly, fledging success rate also increased with clutch size, suggesting that the positive effect of reproductive investment extended beyond hatching and enhanced offspring survival through to fledging (*β* ± SE = 0.075 ± 0.035, *t* = 2.182, *P* = 0.031, n = 152; Figure 3b; Table S7, Model 12).

**Figure 3.**
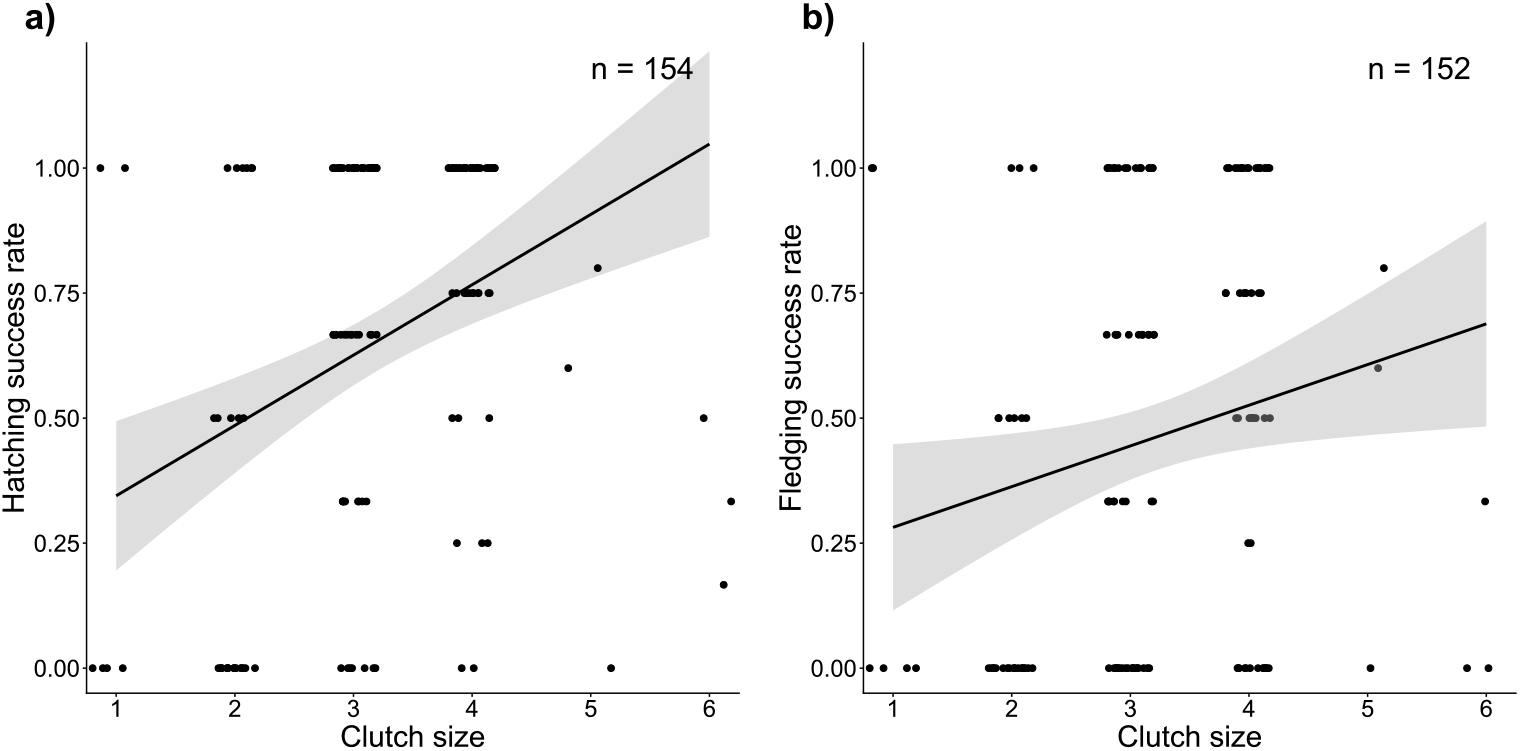
The relationship between the clutch size and the hatching success rate (a, Table S7, Model 11), as well as the fledging success rate (b, Table S7, Model 12). Fitted lines and confidence intervals were obtained using simple linear regression for visualization purposes.

### Effects of nesting type on reproductive success rate

We found no significant effect of nesting type (colonial vs. solitary breeding) on reproductive success in our study. Specifically, hatching success rate did not differ significantly between colonial and solitary nests (*β* ± SE = −0.016 ± 0.065, *t* = −0.251, *P* = 0.802, n = 154; Table 2, Model 13). Similarly, fledging success rate was not significantly associated with nesting type (*β* ± SE = 0.028 ± 0.068, *t* = 0.411, *P* = 0.682, n = 152; Table 2, Model 14).

**Table 2.**
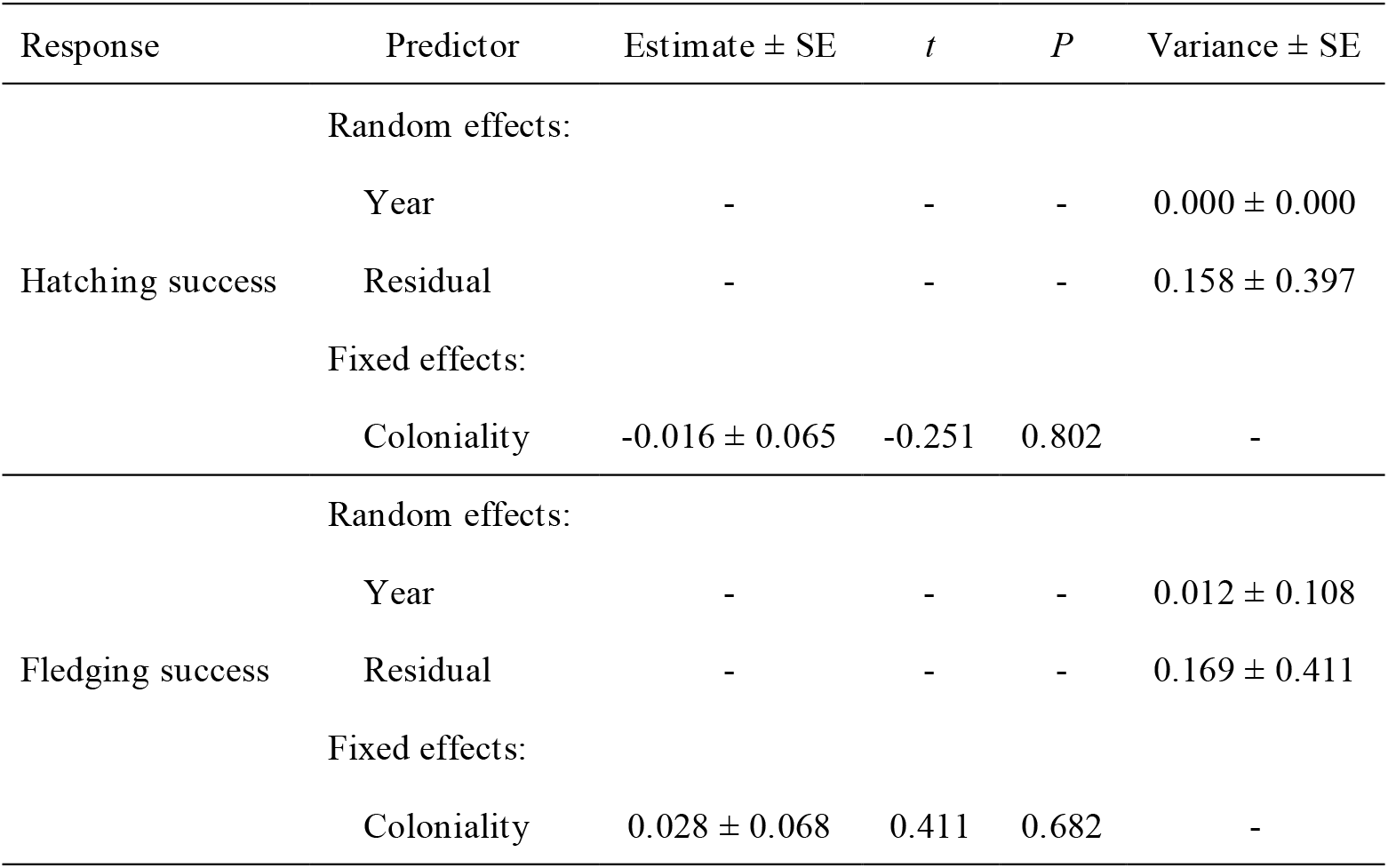
Effects of nesting type on hatching and fledging success rate based on linear mixed-effects models (Model 13 and 14).

## Discussion

In this study, we provide empirical evidence for the effects of breeding timing, clutch size, and breeding strategy (colonial versus solitary nesting) on reproductive success in crested ibis, an endangered and flagship species. These patterns were consistent across the three breeding seasons we examined: reproductive success declined with delayed reproduction, increased with larger clutch sizes, and was not significantly influenced by whether the breeding pairs nested colonially or solitarily. Studying the reproductive biology of endangered species is challenging because it ideally requires intensive monitoring of large numbers of individuals with long-term datasets. Understanding the reproductive performance and its influencing factors is of great importance because this knowledge can help evaluate the populations’ conservation status and predict its evolutionary potential. This is particularly true for flagship or umbrella species that enhance public engagement and promote the protection of broader ecosystems. In this case, our findings offer a comprehensive perspective on how key life-history factors shape reproductive outcomes in wild crested ibis populations and provide a scientific foundation for evaluating the current status of population recovery and refining conservation strategies.

During our study period, except for 2024, when fledging success was significantly higher than in 2023, reproductive success did not differ significantly among the three years, suggesting that interannual environmental conditions did not have a marked effect on crested ibis reproduction. Moreover, the effects of breeding timing, clutch size, and nesting strategy on reproductive success were consistent across the three years, suggesting that our results are robust and repeatable. Nevertheless, previous studies have suggested that long-term climate change and phenological mismatches caused by extreme weather events may influence avian reproductive success, as well as the relationship between life-history traits and reproductive outcomes (Bowers et al., 2016). In a 25-year long-term study of the greater snow goose (*Anser caerulescens atlantica*), researchers found that climate warming advanced plant phenology, and increased the mismatch between the timing of nestling hatch and the peak nutritive quality of plants. As the optimal laying date yielding the highest reproductive success occurred earlier over time, the seasonal decline in reproductive success became more pronounced (Reséndiz-Infante & Gauthier, 2020). Similarly, a long-term population study of the pied flycatcher (*Ficedula hypoleuca*) demonstrated that when the food peak advanced due to increasing temperatures, the decline in clutch size with later egg-laying dates became steeper. (Both & Visser, 2005). Therefore, continued monitoring is essential to further test these hypotheses and provide a reliable data foundation for predicting population reproduction and adaptive responses under rapid climate change (Lack, 1964).

Our study found that both hatching success and fledging success of the crested ibis declined with delayed breeding. As documented in previous studies, multiple mechanisms may contribute to such seasonal declines in reproductive success. One potential explanation is the progressive deterioration of environmental conditions over the breeding season. For instance, in the scarlet ibis (*Eudocimus ruber*), pairs initiating reproduction during later breeding pulses often experience substantially lower reproductive success due to increased frequency of adverse weather events, such as storms (Martínez & Rodrigues, 1999; Olmos, 2003). Similarly, in our study site, Dongzhai, the onset of summer is frequently accompanied by heavy rainfall and strong winds. These conditions can reduce food availability, damage nests, or induce abrupt temperature fluctuations, all of which may negatively affect reproductive performance. Comparable patterns have also been reported in populations from Shaanxi Province, where lower fledging rates were associated with higher rainfall and lower temperatures in May (Yu et al., 2006; Zhai et al., 2011). In addition, prolonged high temperatures following sudden extreme weather events later in the breeding season may interact with parasitic infections to increase individual mortality risk. Parasitism is widespread in crested ibis, but infections are generally non-lethal and only become fatal when individuals are simultaneously exposed to adverse conditions such as food scarcity or extreme weather (e.g., lasting heat wave) (Liu & Yu, 1999; Ding, 2004; Yu et al., 2006; Xi et al., 2025). This effect is likely due to increased energetic costs of parasitic infection, which may reduce the individual’s capacity to cope with environmental stressors. Another potential effect involves increasing predation pressure as the breeding season progresses. Rising temperatures may enhance the activity of predators, such as the king ratsnake (*Elaphe carinata*), thereby elevating predation risk for eggs and nestlings and further reducing reproductive success (Zhang et al., 1999; Jiang et al., 2023).

Beyond environmental, parasitism, and predation-related factors, trade-offs in parental investment are also likely to play a role. As the breeding season progresses, parents must allocate limited energy between current reproduction and subsequent life-history demands, including molt, migration, overwinter survival, and future reproduction (Siikamäki, 1998; Schindler et al., 2024). Greater allocation to self-maintenance later in the season may constrain investment in eggs or offspring, ultimately reducing reproductive success. Empirical support for such trade-offs has been reported in a semi-free-ranging population of crested ibis, where breeding date was negatively associated with both fitness and molt quality: early breeders exhibited higher reproductive output and superior plumage quality, whereas late breeders performed poorly, indicating a clear trade-off between breeding and molt (Li et al., 2025). In addition, individual heterogeneity may further contribute to seasonal declines in reproductive success. According to the ‘quality’ hypothesis, individuals of higher phenotypic or physiological quality tend to breed earlier and achieve greater reproductive success (Gustafsson et al., 1994; Liu et al., 2025), which can generate a negative relationship between breeding date and reproductive success. However, our study was constrained by incomplete individual identification and the lack of experimental manipulation, which precluded a direct assessment of parental quality. Thus, further comprehensive long-term monitoring is required to disentangle these interacting factors.

Our study also revealed that reproductive success in the crested ibis increased with clutch size. Previous studies have declared that clutch size can be associated with breeding dates (Crick et al., 1993), reflecting trade-offs between reproduction and parental survival. Late breeders may reduce clutch size either to advance hatching dates or to limit reproductive investment late in the season, thereby mitigating fitness costs associated with delayed breeding (Siikamäki, 1998; Bêty et al., 2003). In addition, consistent with the ‘quality’ hypothesis, individuals in better condition may be less impacted by adverse environmental conditions and tend to produce both larger and earlier clutches (Rowe et al., 1994; Jean-Gagnon et al., 2018). Consequently, larger clutch size can be associated with higher reproductive success. In our study, we found no significant correlation between clutch size and breeding time (LMM: *β* ± SE = −0.014 ± 0.008, *t* = −1.70, *P* = 0.091), suggesting that clutch size itself may exert an independent effect on reproductive success. From an environmental perspective, greater food availability can simultaneously promote larger clutch sizes and enhance parents’ capacity to successfully rear more offspring (Lack, 1947; Martin, 1987). Consistent with this interpretation, previous studies have shown that food supplementation increases foraging success, clutch size, and the number of fledglings in crested ibis (Lu, 1989; Ding, 2004; Yu et al., 2006). Given that the species primarily feeds on small vertebrates and invertebrates inhabiting wetlands and paddy fields (Ding, 2004; Wu, 2019), and that Dongzhai National Nature Reserve is characterized by extensive wetlands, abundant rice paddies, and high prey diversity (Wu, 2019), the positive relationship between clutch size and reproductive success observed in this population may be partly attributable to favorable environmental conditions and abundant food resources in this region.

Meanwhile, a larger clutch itself may better buffer against offspring loss from predators (Olmos, 2003), as the loss of a single egg or chick represents a smaller proportional reduction in broods containing three eggs than those with only two. This buffering effect may be further reinforced by the pronounced asynchronous hatching characteristic of the crested ibis (Li & Huang, 1986; Shi & Yu, 1989; Zhai et al., 2011). A certain degree of developmental differences arising from asynchronous hatching can enhance reproductive success. It may stagger peaks in begging intensity and food demand, allowing parents to allocate provisioning effort more efficiently across offspring (Fujioka, 1985). When environmental conditions are unfavorable, such hierarchies may also facilitate early brood reduction, whereby younger and weaker nestlings are eliminated earlier, thereby reducing ineffective parental investment and increasing overall reproductive efficiency (Hahn, 1981). An experiment with cattle egret (*Bubulcus ibis*) showed that broods with no hatching intervals and doubled intervals, created by chicks exchanging between nests, had lower parental care efficiency than broods with normal intervals. Synchronized hatching increased aggressive interactions and chick mortality, whereas excessive asynchrony amplified competitive asymmetries, increasing starvation risk for the youngest nestlings (Mock & Ploger, 1987). Thus, under natural conditions, asynchronous hatching in multiple-egg clutches may enhance reproductive success by providing flexibility in brood reduction and parental allocation. More importantly, although sibling competition during feeding does occur in the crested ibis, it appears to be largely facultative rather than obligate, and is considerably less intense than that observed in other closely related species within the family Threskiornithidae. Older and stronger nestlings often relinquish feeding positions after satisfying immediate nutritional needs, rather than persistently suppressing the weaker ones, and at least half of all pecks were false pecks, in which the chicks did not make contact with their siblings even when within reach (Li et al., 2004). In contrast, in the closely related great egret (*Casmerodius albus*), chicks fight frequently regardless of food levels and can result in death (Mock & Parker, 1986; Mock et al., 1990). Such interspecific differences in the severity of sibling competition may buffer intra-brood competition against increases in clutch size, which could help explain the relatively rapid population recovery observed in the crested ibis. Collectively, the buffering effect of larger clutches, the adaptive consequences of asynchronous hatching, and the predominantly facultative nature of sibling competition provide a plausible mechanistic explanation for the strong positive association between clutch size and reproductive success observed in the Dongzhai population.

As expected, neither solitary nor colonial nesting had a significant effect on hatching success or fledging success of the crested ibis in Dongzhai. In our study, the proportion of solitary to colonial nesting was approximately 3:2, indicating a relatively small difference in frequency between these two strategies. This pattern may suggest that current environmental conditions do not strongly favor or constrain either nesting strategy. Such a balance is consistent with the coexistence of opposing selective pressures. On the one hand, abundant resources and predation pressure from snakes and raptors may favor colonial nesting (Yu et al., 2006, 2015; Xi et al., 2025); on the other hand, relatively high parasitic infection rates in this species may promote solitary nesting as a means of reducing parasite transmission (Liu & Yu, 1999; Ding, 2004). Moreover, previous studies have shown that the crested ibis can adopt different breeding strategies over time and across habitats (Shi & Cao, 2001; Liu et al., 2020), highlighting its capacity to flexibly adjust breeding behavior in response to local environmental conditions, such as variation in resource availability. Consequently, the coexistence of all contrasting pressures, together with the species’ inherent capacity for flexible behavioral responses, may therefore maintain both nesting strategies within the population. However, due to limitations in individual identification, information on the genetic relationships among breeding individuals is currently lacking, which prevents us from assessing the influence of kinship on breeding strategies and calls for further comprehensive investigation.

Compared with previous studies, our research not only documents the overall breeding ecology of the crested ibis in Dongzhai across three consecutive breeding seasons, but also, for the first time, quantitatively assesses the effects of breeding timing, clutch size, and nesting strategy (solitary versus colonial) on reproductive success. These findings provide a more integrative understanding of the breeding ecology of the crested ibis and extend existing knowledge of the factors shaping reproductive performance in this endangered species. Nevertheless, it should be noted that constraints in capturing and sampling wild breeding adults currently prevent complete identification of all breeding individuals, limiting our ability to explicitly account for the effects of inbreeding, kinship, and parental quality. These unmeasured factors may have introduced unquantified variation into our analyses. Previous studies have demonstrated that inbreeding can negatively affect embryo survival and individual fitness (Fu et al., 2019; Zheng et al., 2024), and that breeding time as well as clutch size are associated with parental quality and/or territory characteristics (Daan et al., 1989; Siikamäki, 1995; Bêty et al., 2003; Liu et al., 2025). Incorporating these effects will therefore be an important goal for future research. In recent years, we have initiated systematic banding and biological sampling of newly fledged individuals, many of which are now gradually recruiting into the breeding population. We believe that with continued accumulation of marked individuals over time, this effort will enable more detailed, individual-based, and longitudinal monitoring of the Dongzhai population, thereby allowing future studies to explicitly integrate genetic, demographic, and phenotypic factors. Ultimately, a deeper understanding of the factors influencing reproductive success in the crested Ibis will improve our ability to assess the current trajectory of population recovery and provide valuable empirical evidence to develop more targeted conservation strategies and reconcile species conservation with local socio-economic development.

## Conclusion

Our study demonstrates that, for the wild-reintroduced crested ibis population in Dongzhai, reproductive success declined with delayed breeding, increased with larger clutch sizes, and was not significantly influenced by whether the breeding pairs nested colonially or solitarily across three consecutive breeding seasons. Our findings contribute to a more integrative understanding of the breeding ecology of the crested ibis and provide empirical support for improving conservation and management strategies during ongoing population recovery. Although current limitations in individual identification preclude explicit evaluation of genetic and parental-quality effects, ongoing banding and sampling efforts will enable more refined, individual-based analyses in the future.

