## Supplementary material for "The effects of breeding timing, clutch size, and nesting strategy on reproductive success in the crested ibis (*Nipponia nippon*)": Table S1

Table S1. The frequencies of different breeding colony sizes.

| Colony size  (Number of breeding pairs) | Number of colonies |
| --- | --- |
| 2 | 11 |
| 3 | 4 |
| 4 | 1 |
| 5 | 2 |
| 6 | 1 |
| 8 | 1 |
| 10 | 1 |
