## Supplementary material for "The effects of breeding timing, clutch size, and nesting strategy on reproductive success in the crested ibis (*Nipponia nippon*)": Table S2

Table S2: Summary of sample size for this study during 2023–2025.

| Year | 2023 | 2024 | 2025 | Overall |
| --- | --- | --- | --- | --- |
| Number of attempts | 45 | 64 | 67 | 176 |
| Number of nests | 41 | 58 | 61 | 160 |
| Number of eggs | 126 | 187 | 189 | 502 |
| Number of hatched eggs | 72 | 135 | 133 | 340 |
| Number of fledglings | 37 | 109 | 90 | 236 |
