## Supplementary material for "The effects of breeding timing, clutch size, and nesting strategy on reproductive success in the crested ibis (*Nipponia nippon*)": Table S3

Table S3. The effect of breeding timing on hatching success rate based on the linear mixed-effects model (Model 3).

|  | Estimate ± SE | *t* | *P* | Variance ± SE |
| --- | --- | --- | --- | --- |
| Random effects: |  |  |  |  |
| Year | - | - | - | 0.000 ± 0.007 |
| Residual | - | - | - | 0.122 ± 0.349 |
| Fixed effects: |  |  |  |  |
| First egg-laying date | -0.016 ± 0.003 | -5.217 | **< 0.001** | **-** |
