## Supplementary material for "The effects of breeding timing, clutch size, and nesting strategy on reproductive success in the crested ibis (*Nipponia nippon*)": Table S4

Table S4. Year-specific effects of breeding timing on hatching success rate based on linear models for 2023–2025 (Model 5–7).

|  | Predictor | Estimate ± SE | *t* | *P* |
| --- | --- | --- | --- | --- |
| 2023 | Intercept | 0.879 ± 0.119 | 7.383 | - |
|  | First egg-laying date | -0.012 ± 0.005 | -2.26 | **0.030** |
| 2024 | Intercept | 1.130 ± 0.134 | 8.438 | - |
|  | First egg-laying date | -0.021 ± 0.006 | -3.369 | **0.001** |
| 2025 | Intercept | 0.977 ± 0.096 | 10.22 | - |
|  | First egg-laying date | -0.019 ± 0.005 | -3.572 | **< 0.001** |
