## Supplementary material for "The effects of breeding timing, clutch size, and nesting strategy on reproductive success in the crested ibis (*Nipponia nippon*)": Table S5

Table S5. The effect of breeding timing on fledging success rate based on the linear mixed-effects model (Model 4).

|  | Estimate ± SE | *t* | *P* | Variance ± SE |
| --- | --- | --- | --- | --- |
| Random effects: |  |  |  |  |
| Year | - | - | - | 0.012 ± 0.111 |
| Residual | - | - | - | 0.146 ± 0.382 |
| Fixed effects: |  |  |  |  |
| First egg-laying date | -0.015 ± 0.003 | -4.380 | **< 0.001** | **-** |
