## Supplementary material for "The effects of breeding timing, clutch size, and nesting strategy on reproductive success in the crested ibis (*Nipponia nippon*)": Table S6

Table S6. Year-specific effects of breeding timing on fledging success rate based on linear models for 2023–2025 (Model 8–10).

|  | Predictor | Estimate ± SE | *t* | *P* |
| --- | --- | --- | --- | --- |
| 2023 | Intercept | 0.659 ± 0.121 | 5.425 | - |
|  | First egg-laying date | -0.016 ± 0.005 | -2.980 | **0.005** |
| 2024 | Intercept | 0.981 ± 0.157 | 6.237 | - |
|  | First egg-laying date | -0.020 ± 0.007 | -2.647 | **0.011** |
| 2025 | Intercept | 0.672 ± 0.102 | 6.597 | - |
|  | First egg-laying date | -0.012 ± 0.006 | -2.189 | **0.033** |
