## Supplementary material for "The effects of breeding timing, clutch size, and nesting strategy on reproductive success in the crested ibis (*Nipponia nippon*)": Table S7

Table S7. Effects of clutch size on hatching and fledging success rates based on linear mixed-effects models (Model 11 and 12).

| Response | Predictor | Estimate ± SE | *t* | *P* | Variance ± SE |
| --- | --- | --- | --- | --- | --- |
| Hatching success rate | Random effects: |  |  |  |  |
|  | Year | - | - | - | 0.000 ± 0.000 |
|  | Residual | - | - | - | 0.140 ± 0.374 |
|  | Fixed effects: |  |  |  |  |
|  | Clutch size | 0.141 ± 0.032 | 4.447 | **< 0.001** | **-** |
| Fledging success rate | Random effects: |  |  |  |  |
|  | Year | - | - | - | 0.010 ± 0.100 |
|  | Residual | - | - | - | 0.164 ± 0.405 |
|  | Fixed effects: |  |  |  |  |
|  | Clutch size | 0.075 ± 0.035 | 2.182 | **0.031** | **-** |
